# Origin of the far-red absorbance in eustigmatophyte algae red-shifted Violaxanthin-Chlorophyll *a* Protein

**DOI:** 10.1101/2024.04.27.591434

**Authors:** Alessandro Agostini, David Bína, Dovilė Barcytė, Marco Bortolus, Marek Eliáš, Donatella Carbonera, Radek Litvín

## Abstract

Photosynthetic organisms harvest light for energy. Some eukaryotic algae have specialized in harvesting far-red light by tuning chlorophyll *a* absorption through a mechanism still to be elucidated. Here, we combined optically detected magnetic resonance and pulsed electron paramagnetic resonance measurements on red-adapted light-harvesting complexes, rVCP, isolated from the freshwater eustigmatophyte alga *Trachydiscus minutus* to identify the location of the pigments responsible for this remarkable adaptation. The pigments have been found to belong to an excitonic cluster of chlorophylls *a* at the core of the complex, close to the central carotenoids in L1/L2 sites. A pair of structural features of the Chl *a*403/*a*603 binding site, namely the histidine-to-asparagine substitution in the magnesium-ligation residue and the small size of the amino acid at the *i*-4 position, are proposed to be the origin of this trait. Phylogenetic analysis of various eukaryotic red antennae identified several potential LHCs that could share this tuning mechanism.

## Introduction

Photosynthetic eukaryotes rely mainly on the proteins of the light-harvesting complex family (LHCs) to perform the important role of harvesting and transferring energy to the photosynthetic reaction centres. LHCs are characterised by a conserved structural blueprint consisting of three transmembrane α-helices binding chlorophylls (Chls) and carotenoids (Cars)^1^. An essential part of their architecture consists of a left-handed coiled-coil helix pair constituted by the two homologous α-helices A and B, which generate two conserved carotenoid binding sites, labelled L1 and L2 in the light harvesting complex II (LHCII) of higher plants (Fig. 1a)^2^. An eight-pigment cluster, constituted by these two Cars each strongly interacting with an excitonically-coupled cluster of three Chls (see Fig. 1), has been shown to be pivotal in both the light-harvesting^3,4^ and photoprotective^5–9^ properties of these LHCs. Pigment binding sites of helices A and B are strongly conserved among different LHCs, with modifications limited to substitutions of individual chlorophyll/carotenoids. Pigment binding sites closer to helix C are less conserved^2,10–12^.

**Fig. 1.**
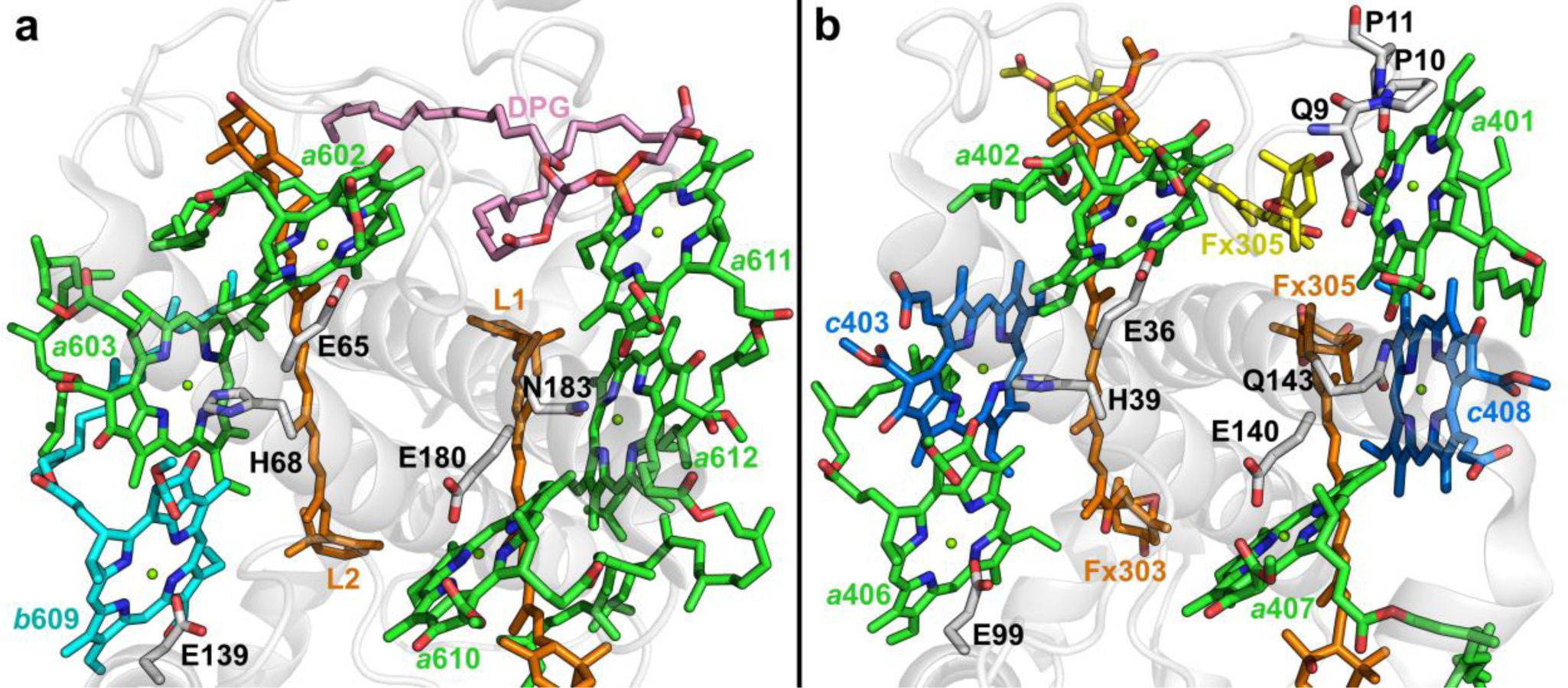
Comparison of prototypical LHCs. Lumenal view of **(a)** *Spinacia oleracea* LHCII (PDB ID: 1RWT^2^) and **(b)** *Phaeodactylum tricornutum* FCP (PDB ID: 6A2W^10^), focussing on the conserved pigments close to the L1/L2 sites. Green sticks, Chls *a*; cyan sticks, Chl *b*; blue sticks, Chls *c*; orange and yellow sticks, luteins and fucoxanthins; pink sticks, dipalmitoyl-phosphatidylglycerol (DPG); white cartoons, polypeptide chain.

One aspect of photosynthetic organisms that holds both evolutionary and technological interest is the ability to adapt to different light conditions. LHC proteins provide flexible scaffolds for binding of diverse pigment mixtures. An example is the fucoxanthin-chlorophyll *a*/*c* protein complex of diatoms (FCP, Fig. 1b), which achieves excellent light-harvesting efficiency in the blue-green part of the spectrum through the utilization of Chl *c* and the carotenoid fucoxanthin. This is optimised for survival in the water column, where the red part of the spectrum is strongly depleted through absorption of water^13^. This approach has been also emulated *in vitro* by attaching artificial pigments to light-harvesting complexes to enhance their absorption in the green gap^14,15^.

Another adaptation to light limitation is the expansion of the chlorophyll absorption to the far-red part of the light spectrum. This has been most studied in cyanobacteria synthesising specific low-energy chlorophyll species, Chl *d*^16^ and Chl *f*^17^. While these pigments are specifically cyanobacterial, Chl *a*-to-Chl *d* substitution has been performed in LHCII producing a red-shifted LHC complex^18^. The promise of this research direction lies in harvesting the portion of light most available in dense canopies^19,20^.

Under natural conditions, eukaryotes achieve the red-shift of LHC absorbance without the need to produce specialized pigments by relying on pigment-pigment and pigment-protein interactions^21–29^.

Some of the redmost LHCs have been identified in algae belonging to the class Eustigmatophyceae^21,24–26^. Eustigmatophyte LHCs bind solely Chl *a* and carotenoids violaxanthin and vaucheriaxanthin, and are therefore called Violaxanthin-Chlorophyll *a* Proteins (VCP)^30–33^. Red-shifted VCP (rVCP) has been so far isolated from two eustigmatophyte species^24,25^ and it has been speculated that the red-shifted absorption is the result of an excitonic coupling of pigments located on the interface between subunits of an oligomeric LHC complex^34^.

In the present study, we employed photogenerated triplet states as internal probes in a magneto-optical spectroscopic investigation that, along with new knowledge of the rVCP peptide sequences of the eustigmatophyte *Trachydiscus minutus* (*Tm*), allowed us to distinguish and localise the different Chl pools, aiming to understand the mechanism of the colour tuning in rVCP.

## Results

### Pigment and protein composition of rVCP in *T. minutus*

rVCP binds 19 violaxanthin and 10 vaucheriaxanthin per 100 Chl *a*^34^, differing from VCP in the almost complete lack of esterified vaucheriaxanthin (10% of the vaucheriaxanthin pool) and in the pigment ratio (3.4: 1 Chl *a*: Car) that is more similar to plant LHCII^2^ than VCP^30,31,34^.

The composition of purified *Tm* rVCP protein was analysed using tandem mass spectrometry (MS/MS): the sample was predominantly composed of three polypeptides (see Supplementary Table S1) referred to as DN2982, DN29098, and DN6201 (following the IDs of the sequence contigs in the transcriptome assembly). The rVCP polypeptide sequences were aligned against antenna sequences from the diatom *Phaeodactylum tricornutum* (*Pt*) *i.e.* Lhcf4, the canonical FCP protein, and Lhcf15, the building block of the diatom red-shifted antenna^35–38^ (Fig. 2).

**Fig. 2.**
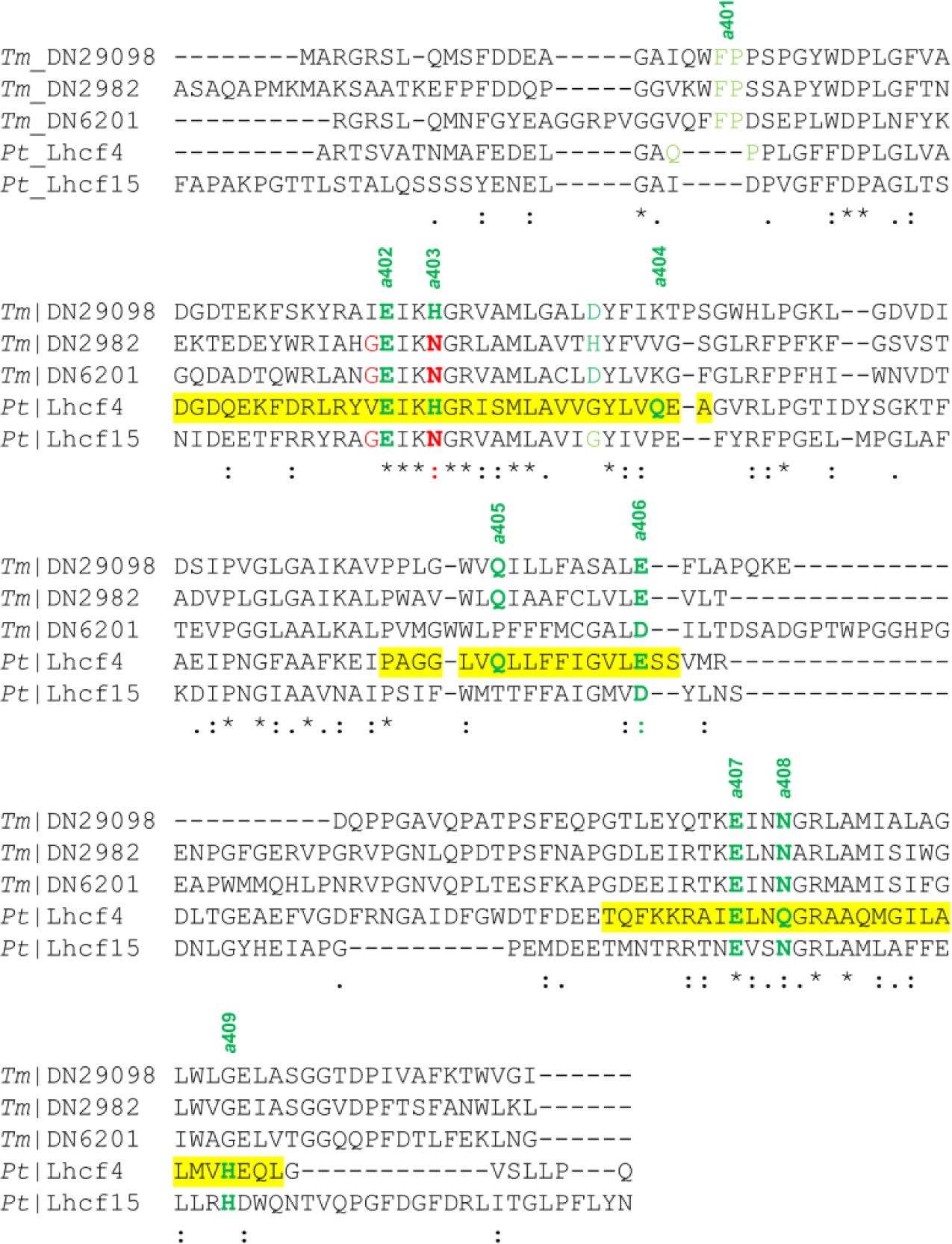
Alignment of polypeptide sequences of LHCs. Polypeptide sequences of the LHCs from *Trachydiscus minutus* (*Tm*) and *Phaeodactylum tricornutum* (*Pt*)^44,45^ were used for the alignment. Yellow background signifies a transmembrane helix (derived from the crystallographic structures 6A2W^10^). Chl binding residues in the sequences are color-coded: green, Chl side chain ligand; light green, Chl backbone ligand via a water molecule; red, Chl red-shifting GxxxN binding motif. Note that in the case of the *a*404 site, a suitable ligand is found at the *i*-4 position in the *Tm* sequences (similarly to the binding of *a*604 in LHCII^2^), which is proposed to replace the ligation in *Pt*.

The sequence alignment revealed a clear conservation of the amino acid residues involved in the binding of several pigments (see Figs. 2 and S1 for Chls and Cars, respectively). Five Chls *a* (namely, *a*402, *a*403, *a*406, *a*407, and *a*408) appear strictly conserved in rVCP (we will refer to the Chl-binding sites named *c*403 and *c*408 by Wang et al.^10^ as *a*403 and *a*408, since rVCP binds only Chl *a*^34^). Note that in two of the three *Tm* rVCP sequences, the ligand of the Chl *a*403 site is an asparagine (Asn), substituting the histidine (His) commonly present at this site^2,10^, and Asn in this position was connected to the red-shifting of Lhca3-4 in plants^27,39,40^.

The *a*405 site seems to be retained in two of the three *Tm* sequences (DN2982 and DN29098). Regarding the *a*404 site, a direct binding as observed in *Pt* FCP can be excluded, but direct binding from the residue at the position *i*-4, similarly to *Gephyrocapsa* (=*Emiliania*) *huxleyi E-*FCP^41^, can be proposed. The retainment of *a*401 is difficult to assess due to its binding mode: a phosphoryl group of a lipid in the case of LHCII, and a QPP motif in a loop in the case of FCPs^10,42^. A binding mode like the latter can be proposed to take place also in rVCPs, due to the conservation of an FP pair of amino acid residues at a similar position.

Some of the Chl binding sites discussed above (*a*401, *a*404, *a*405, *a*409) are also probably conserved, since the binding of at least 7 Chls *a* and 2 Cars is expected for rVCP from the stoichiometries determined via HPLC and the *Pt* FCP structure analogy^10,34^. Carotenoid binding sites L1 and L2 are definitely conserved (see Supplementary Fig. S1), congruently with their known structural role in LHCs^43^.

The higher sequence similarity of the three *Tm* sequences with *Pt* Lhcf4 (21-26 %), when compared to the one with LHCII (16-17 %), suggests the use of the *Pt* FCP structure as a more adequate structural model for rVCP in the following analyses.

### Identification of the site energy of the unquenched ^3^Chls *a*

Illumination of rVCP at cryogenic temperatures leads to the formation of ^3^Chl *a,* which can be detected by Fluorescence Detected Magnetic Resonance (FDMR). The transitions between the triplet sublevels of the ^3^Chl *a* (namely, |D|−|E| and |D|+|E|, see Fig. 3a) are detected by monitoring the emission of the sample as a function of the frequency of a microwave radiation (650-1050 MHz)^46^. The presence of unquenched ^3^Chl *a* is a common finding in LHCs despite the presence of photoprotective carotenoids^5,47–49^ and has been previously reported also for VCP^5,47^.

**Fig. 3.**
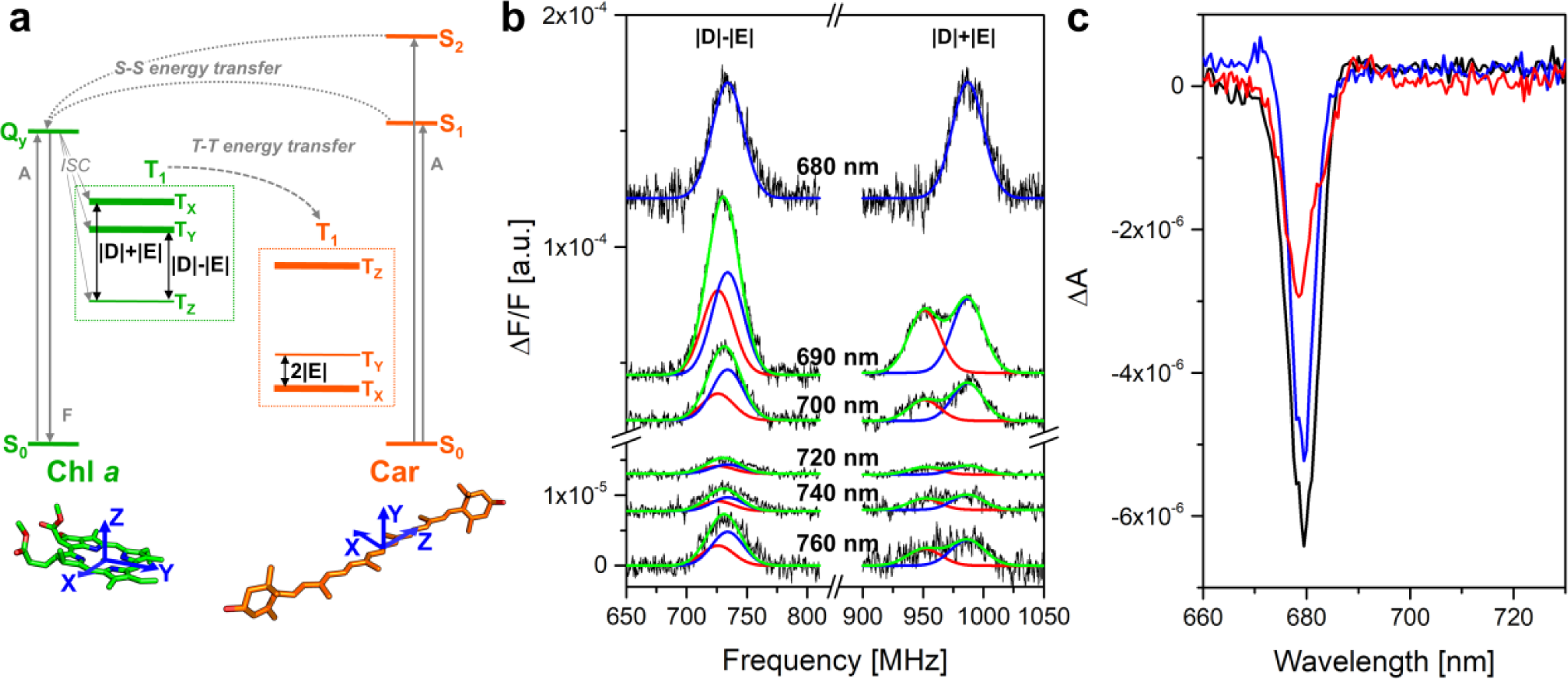
^3^Chl *a* optically detected magnetic resonance (ODMR) spectra of rVCP. **(a)** Jablonski diagram of the main electronic states of Chls and Cars in LHCs (in green and orange, respectively). The states are arranged vertically by energy (not in scale), and horizontally by spin multiplicity. Absorption (A), fluorescence (F), and ISC are indicated by straight grey arrows; singlet-singlet (S-S) and triplet-triplet (T-T) energy transfers by curved dashed grey arrows; the transitions between the spin sublevels by double-pointed black arrows. For readability, only the transitions more relevant for the discussion of the results are drawn. The triplet sublevels are highlighted by dashed boxes for the two molecules. The relative populations of the triplet sublevels are indicated by the thickness of the level bars. At the bottom of the panel, the molecular structure of chlorophyll *a* (Chl *a*) and lutein (Car) with the directions of the zero-field splitting principal axes (zfs, the axes of the dipolar interaction between the two unpaired electrons) are reported (in blue arrows). **(b)** FDMR spectra (black lines) of the ^3^Chl |D|-|E| and |D|+|E| transitions at different wavelengths in the 680–760 nm range, as indicated. Amplitude modulation frequency 33 Hz, time constant 100 ms, temperature 1.8 K. The spectra are vertically shifted for better comparison. Reconstruction (green lines) of the experimental spectra as a sum of Gaussian components (blue and red lines). The fitting parameters are reported in Supplementary Table S2. **(c)** T-S spectra of ^3^Chl *a*. Resonance frequencies: 733 MHz (black line), 945 MHz (red line), and 1000 MHz (blue line). Amplitude modulation 33 Hz, time constant 1 s, temperature 1.8 K.

The optical properties of rVCP Chls *a* are dominated by a pool of red-shifted Chls, which contribute to a third of the integrated area of the Q_y_ 0-0 absorption band (Fig. S2), and are the prevalent emitters when it comes to fluorescence, particularly at cryogenic temperatures^24^. Therefore, it is remarkable that the largest intensities are found at bluer (680-700 nm) wavelengths (Fig. 3b) when the wavelength-dependence of the ^3^Chl *a* FDMR spectra is analysed. Strong suppression of the ΔF/F ratio for wavelengths higher than 700 nm is an indication that the unquenched ^3^Chls *a* are not significantly connected to the redmost Chls *a*. Two ^3^Chl *a* populations contribute to the FDMR spectra at all wavelengths, except at 680 nm. The two contributions are well resolved in the |D|+|E| transitions and can be disentangled due to their different wavelength dependence (see Fig. 3b).

Further data on the pool of unquenched rVCP ^3^Chl have been collected by an ODMR variant, absorption detected magnetic resonance, by monitoring microwave-induced changes in the sample absorption. The spectra (Triplet minus Singlet, T-S) result from a wavelength sweep while fixing the microwaves at a frequency in resonance with a zero-field splitting (zfs) magnetic transition of ^3^Chl (either |D|-|E| or |D|+|E|, see Fig. 3a). In good agreement with the ^3^Chl FDMR results, the ^3^Chl T-S signal is also arising from the bluer pools of Chls, as shown by the narrow bleaching peaking at 680 nm (Fig. 3c), corresponding to the Q_y_ 0-0 absorption band of the ^3^Chls, and the flat profile for wavelengths longer than 690 nm, a clear indication of a lack of interaction of the unquenched ^3^Chls with redmost chlorophylls.

The absence of signals attributable to triplet states localised on the redmost Chls *a,* which dominate the fluorescence spectra and are expected to be the terminal collectors of the excitation in the antenna system, implies an efficient triplet quenching mechanism on these Chls provided by nearby carotenoids. To characterise these photoprotective pathways, analogous ^3^Car ODMR experiments were performed.

### 3Car ODMR reveals that the redmost Chls are effectively photoprotected

Although carotenoids are non-fluorescent molecules, their FDMR transitions can be indirectly detected via the emission of Chls coupled by energy transfer pathways^46^. The change of the steady state population of the ^3^Car levels, induced by a resonant microwave field, is detected as an intensity change in the fluorescence of nearby coupled Chl^46^. Therefore, the wavelength dependence of ^3^Car FDMR signals can be utilised to selectively investigate the optical properties of just the coupled Chls.

Fig. 4a shows the FDMR spectra detected irradiating around the main resonance transition of ^3^Cars^46^ (2|E|, 210-250 MHz), at various emission wavelengths^24^. The lack of signal in the 690-700 nm region points to a weak or absent coupling of the triplet-carrying carotenoid to the bluer chorophylls, which were found to mostly contribute to the unquenched ^3^Chl pools (seen in Fig. 3b). On the contrary, a strong FDMR signal is found from 710 nm on, a clear indication that ^3^Cars are coupled to a pool of red Chls *a*. The zfs parameters of ^3^Car (|D| = 0.0387 cm^-1^, |E| = 0.0038 cm^-1^) were determined from the 2|E|, |D|-|E| and |D|+|E| transitions at 740 nm. These values are close to the main ^3^Car component observed in VCP (|D| = 0.0393 cm^-1^, |E| = 0.0039 cm^-1^)^5^.

**Fig. 4.**
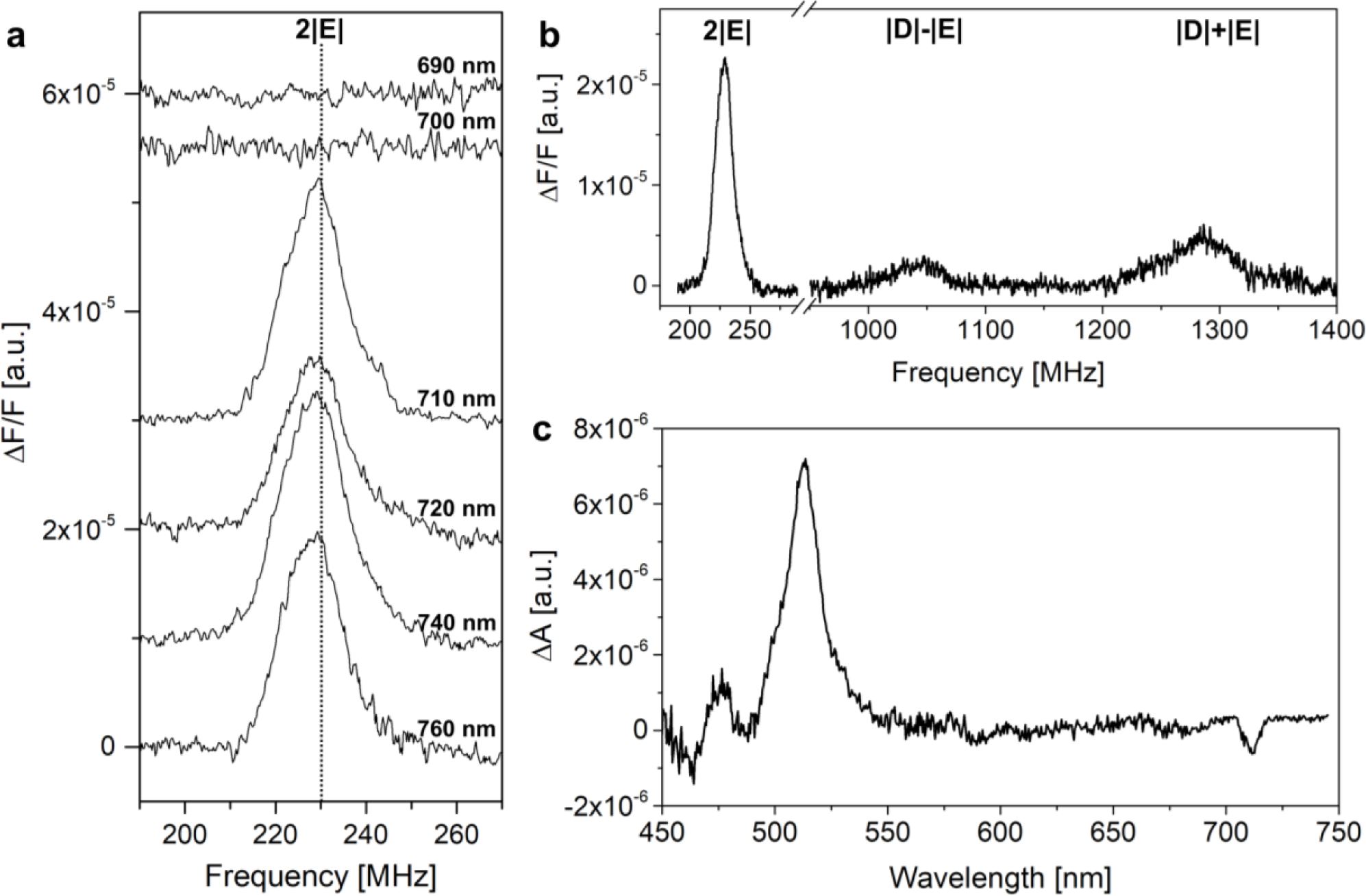
^3^Car ODMR spectra of rVCP. **(a)** FDMR spectra of the ^3^Car 2|E| transition detected at different wavelengths in the 690-760 nm range, as indicated. Amplitude modulation 333 Hz, time constant 100 ms, temperature 1.8 K. The spectra are vertically shifted for better comparison. The vertical dotted line highlights the 2|E| peak position. **(b)** FDMR spectrum of ^3^Car 2|E|, |D|-|E|, and |D|+|E| transitions at 740 nm. Amplitude modulation 333 Hz, time constant 100 ms, temperature 1.8 K. **(c)** T-S spectrum of ^3^Car, obtained with a resonance frequency of 230 MHz (^3^Car 2|E| transition, see panels a-b). Amplitude modulation 333 Hz, time constant 1 s, temperature 1.8 K.

The ^3^Car T-S spectrum (Fig. 4c), obtained with the microwave frequency set at the maximum of the ^3^Car 2|E| transition (230 MHz), is dominated by an intense band corresponding to the triplet-triplet absorption of ^3^Car (512 nm). Its remarkably narrow bandwidth (full width at half maximum, FWHM = 640 cm^-1^) suggests the presence of a single ^3^Car component. The negative signals at 460 nm and 485 nm are due to the bleaching of the carotenoid S_2_ singlet-singlet absorption bands. The T-S spectrum shows a bleaching at 711 nm, i.e. in the region where no carotenoid signals are expected. This, commonly observed in antenna proteins, originates from the electronic coupling of the Car carrying the triplet state with the proximal Chls, as recently explained by Migliore *et al*.^50^. The far-red wavelength of the interaction peak in rVCP is similar to that in plant Lhca4^51^, confirming the strong interaction of the ^3^Car with the redmost pool of Chls *a* in agreement with the ^3^Car FDMR spectra. This Car-Chl *a* interaction peak has a FWHM of 160 cm^-1^, similar to those for the interaction of the luteins in site L1/L2 with the excitonic clusters of *a*610-*a*612-*a*611 and *a*602-*a*603 in plant LHCII^9^. Whereas for Chls *a* with weaker coupling with nearby Chls *a*, such as in dinoflagellate PCP^48^ and LHC^49^, and diatom FCPs^42,52^, the corresponding FHWMs were found to be smaller, in a 90-120 cm^-1^ range.

These results show that far-red absorbing chlorophylls are in close proximity to the carotenoids that populate the triplet state. To determine the mutual orientation of the Chl-Car triplet-triplet energy transfer (TTET) pair identified from the ^3^Car T-S spectrum, we measured the ^3^Car using pulse electron paramagnetic resonance (EPR).

### Identification of the TTET pathways by means of ^3^Car pulse EPR

EPR techniques have proven to be an invaluable asset in obtaining structural information regarding the TTET acceptor-donor pair^6,53^. Since the initial spin polarisation pattern of ^3^Car immediately following TTET is inherited from that of the ^3^Chl donor during TTET and depends on the relative pigment arrangement inside the protein scaffold. The determination of the initial spin polarisation pattern of the ^3^Cars, before the onset of the anisotropic relaxation of its triplet spin sublevels^54,55^, requires a light-induced field-swept electron spin echo (FS-ESE) sequence which selectively suppresses the contribution of ^3^Chl *a*^5,6^, for the strong anisotropic relaxations of porphyrin scaffolds^56^. In light-induced FS-ESE, laser photoexcitation is followed by two nanosecond microwave pulses obtaining an ESE whose integrated intensity is recorded as a function of the intensity of a static magnetic field removing the degeneracy of the triplet sublevels (see Fig. 5a).

**Fig. 5.**
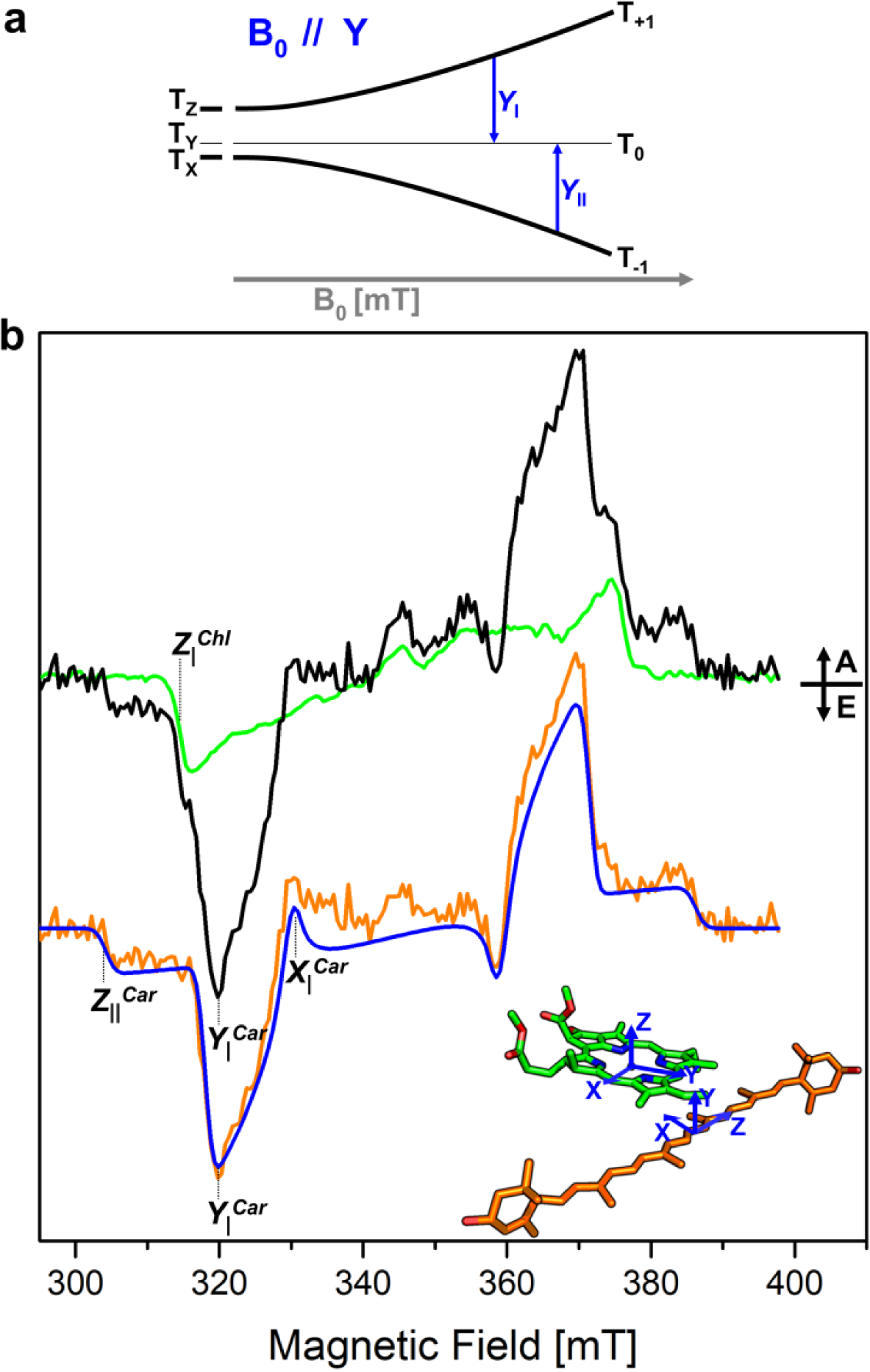
Pulse EPR spectrum of rVCP. **(a)** Scheme of the energies of the triplet sublevels of a ^3^Car (D>0 and E<0) as a function of an external magnetic field, B0, aligned with the ^3^Car zfs axis Y. Whenever the energy of the microwave radiation matches the energy gap between T0 and either T+1 or T-1, a transition can be observed (XI or XII for B0 parallel to Y, respectively). The transitions can be either emissive (E) or absorptive (A) depending on the relative populations of the high-field triplet sublevels involved, indicated by the thickness of the level bars. **(b)** FS-ESE spectrum of rVCP (black) and Chl *a* dissolved in Triton X-100 micelles (green) at 50 K. The difference (orange curve) between the FSE spectrum of rVCP and the FSE spectrum of Chl *a,* which corresponds to the ‘pure’ ^3^Car spectrum, has been vertically translated for clarity. The simulation of the ^3^Car spectrum for Fx303/Fx305 (blue line) is calculated considering a population of the triplet state by means of TTET starting from the triplet state of the closest conserved Chl *a* (Chls *a*403 and *a*408, respectively). The polarizations of the simulated ^3^Car components were determined on the basis of atomic coordinates for the acceptor-donor pairs derived from the crystallographic structure^10^ and an initial donor ^3^Chl polarisation (Px:Py:Pz = 0.375:0.425:0.200)^5^, resulting in a ^3^Car polarization of (Px:Py:Pz = 0.41:0.20:0.39) for both Fx303 and Fx305 (a molecular scheme of the acceptor-donor pair with the zfs tensors of the two molecules is reported at the bottom of the panel). The simulated ^3^Car spectra were calculated using the following parameters: D = -41.0 mT; E = -4.1 mT; linewidths (lwx, lwy, lwz) = (2.0, 2.0, 2.5) mT. Canonical transitions discussed in the text have been highlighted in the low-field half of the spectra. A = absorption, E = emission.

Residual contributions from ^3^Chl *a* were still present in the FS-ESE spectrum of rVCP (Fig. 5b – black), which were subtracted using the FS-ESE of Chl *a* dissolved in Triton X-100 (Fig. 5b – green). The resulting ^3^Car FS-ESE spectrum of rVCP is characterised by an EEAEAA polarisation pattern (Fig. 5b – orange), as already observed in eustigmatophyte VCP^5^ and plant LHCII^6^. When considering the polarizations for the possible couples of Chl and Car, calculated on the basis of atomic coordinates for the acceptor-donor pairs derived from the crystallographic structure^10^, the symmetrically related pairs Chl *a*408-L1 and Chl *a*403-L2 best fitted the experimental spectrum. These polarizations were compatible also with time-resolved EPR spectra reported in the SI (Fig. S3).

The assignment of Chl *a*408-L1 and Chl *a*403-L2 as partners in TTET aligns with these being the closest Chl *a*-Car pairs in terms of π-π and centre-centre distances, both in the LHCII^6^ and in the *Pt* FCP^42^ structures. Together with the finding derived from the analysis of the ^3^Car T-S bleaching at 711 nm (Fig. 4c), we can assign the red excitons to either one or both of the clusters *a*402-*a*403-*a*406 and *a*401-*a*408-*a*407. The present observation that the development of the low-energy states does not entail major changes in pigment organisation from regular LHC agrees with previous conclusions derived from an analysis of the singlet excitation energy transfer dynamics^34^.

## Discussion

A phylogenetic analysis of the antenna sequences unsurprisingly places *Tm* rVCP close to the main antenna proteins of the eustigmatophyte *Nannochloropsis*, Lhcv1/2, within the main group of FCP-like LHC proteins (Fig. S4). This supports earlier proposals that low-energy Chl *a* forms evolved independently in various algal groups and that this evolution requires only minor changes in the protein framework. Focusing on the Chl *a*-binding sites, the multiple alignment of helix B sequences (Fig. 6) shows that in two of the three proteins forming the rVCP complex, the ligand at the *a*403 (*a*603 in plant LHCII) site is an Asn, instead of the typical His. This amino acid exchange is responsible for the development of the red state in plant Lhca4^27,39^. The role of Asn appears to be twofold. Primarily, the smaller volume of the sidechain compared to His residue brings the pigment into closer contact with the pigment bound at the *a*406 (*b*609 in plant LHCII) site. Secondly, as shown recently^40^, the Asn sidechain forms a hydrogen bond to the *a*406 pigment (see Fig. 7), likely stabilising the conformation of the *a*403-*a*406 chlorophyll dimer. The resulting molecular orbital overlap of the paired Chls *a* leads to the development of low-energy states of a mixed excitonic-charge-transfer (CT) nature^40^. In rVCP, the presence of CT states was not demonstrated directly but was inferred from the broadening of the low-temperature emission spectra^24^. One of the three proteins identified in rVCP contains the His-ligand to Chl *a* at the *a*403 position and thus could in principle be lacking the red-Chl *a* forms. However, we detected no Chl *a* absorbing around 680 nm in contact with carotenoids (Fig. 4c). This suggests that the sample consists of heterooligomers, all of which contain the *a*403-Asn protein(s), and hence the excitation energy is always efficiently transferred to the lowest-lying Chl *a* state.

**Fig. 6.**
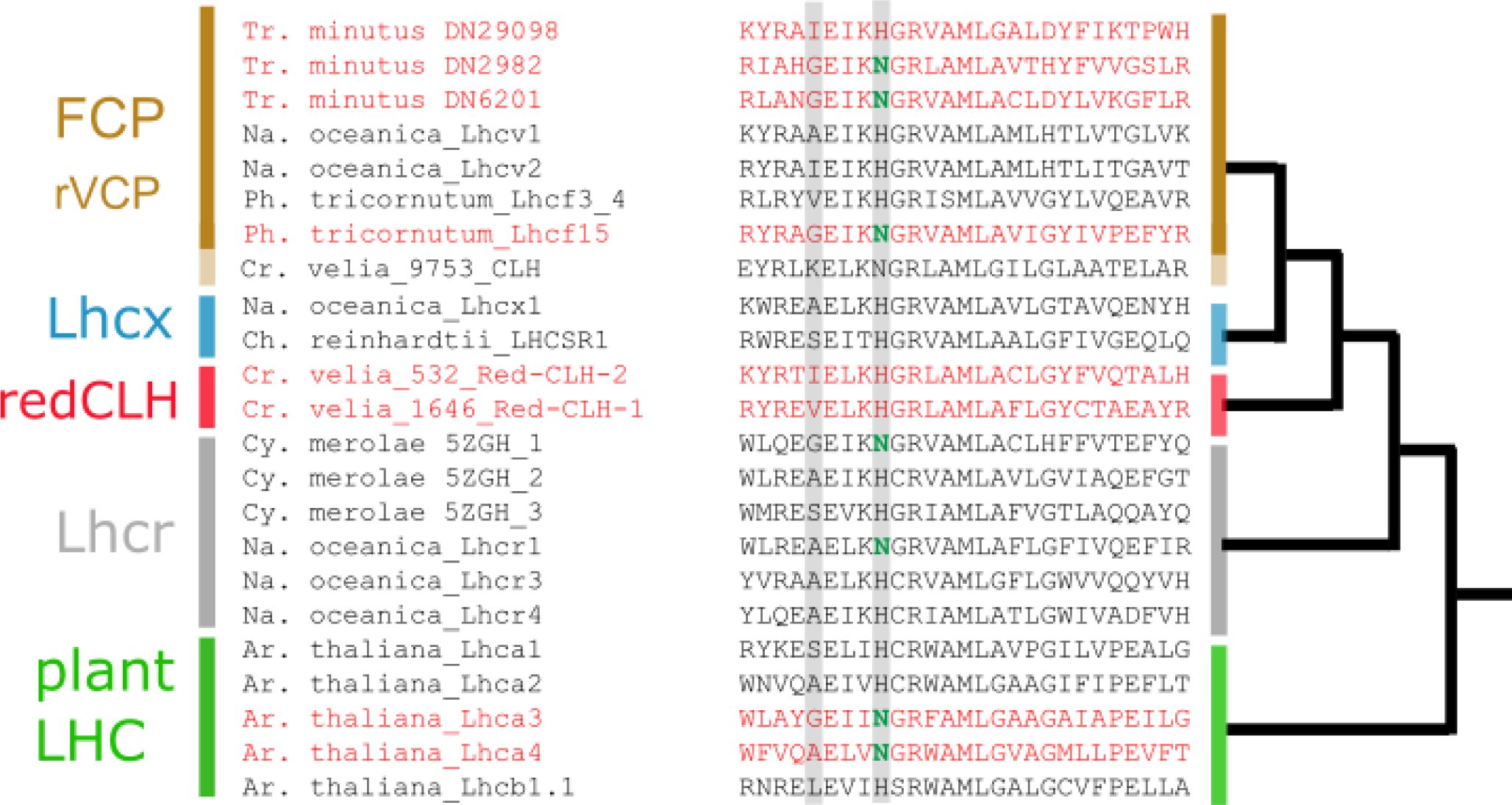
Multiple alignment of helix B sequences of LHC proteins with simplified phylogenetic tree topology on the right. Highlighted is the Chl-binding site *i* = *a*403/*a*603 (H or N) and the corresponding position at *i*-4 (see text for explanation). Detailed version of the phylogenetic tree used to classify the proteins into groups given on the left is presented in Fig. S4. Sequences of proteins that have been purified in a red-shifted Chl *a*-containing form are shown in red.

**Fig. 7.**
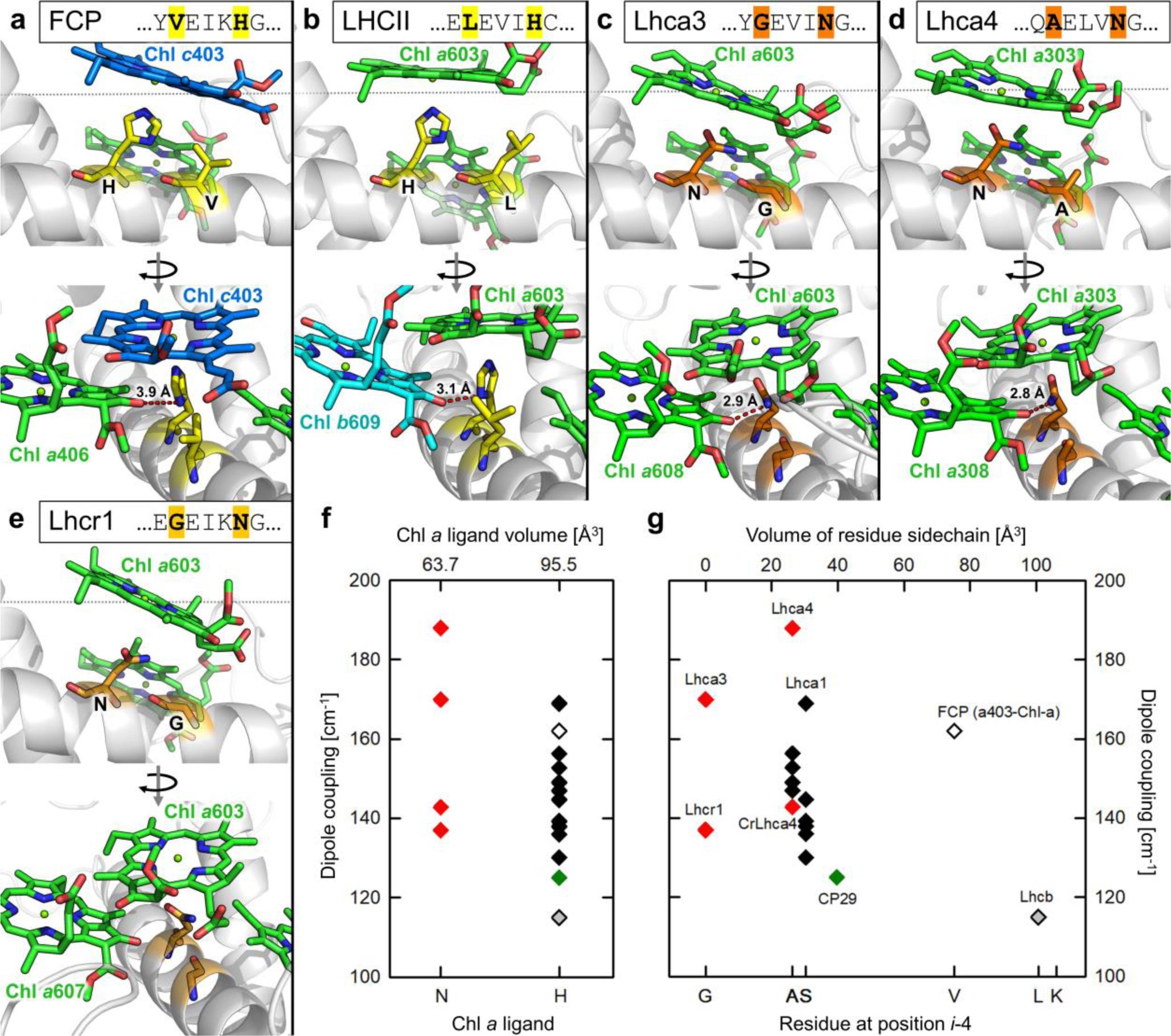
Investigation of the Chl *a*403(*a*603) binding site in the LHC superfamily. (**a-e**) Detailed view of the chlorophyll-binding site *a*403(*a*603) in different LHC proteins (**a**: FCP: pdb 6A2W, **b** LHCII: 1RWT, **c**, **d** Lhca3, 4: 7DKZ, **e** Lhcr1: 5ZGH). (**f**) plot of excitonic coupling between Chl *a* at sites *a*403 and *a*406 according to the Chl *a* ligand (*i* = N or H); (**g**) excitonic coupling vs. the residue at *i*-4 position. Residue sidechain volumes are given on the top horizontal axis. All chlorophylls in the respective sites were parametrized as Chl *a* for the purpose of coupling computation.

The plant photosystem I antenna is so far the only system for which this particular mechanism has been unequivocally demonstrated experimentally and its extrapolation to other systems, while parsimonious, is not granted. Since the *a*403-*a*406 dimer consists of molecules bound to different transmembrane helices of the LHC structure, the coupling ultimately depends on the tertiary structure of the whole protein and could be influenced by factors beyond an exchange of a single amino acid residue. A case in point appears to be the main antenna protein of the alveolate *Chromera* (*C.*) *velia* (19753_CLH), which also contains Asn at the *a*403 position while not exhibiting the spectroscopic signature of a red-shifted chlorophyll^57^. As seen in Fig. 6, a recurring theme of the Chl *a* binding of the red-shifted systems is the presence of a small-sidechain residue (Gly/Ala) in the position *i*-4 from the chlorophyll ligand. Two of the three *Tm* sequences show a conserved glycine at this position, similarly to plant Lhca3 and red algal Lhcr1. As visible from the comparison in Fig. 7a-e, Gly or Ala in position *i*-4 give more space to the Chl in the *a*403 site, so it can be pulled by the shorter Asn ligation without steric clashes between the carbonyl group in position 13^1^ of the isocyclic ring and the side chain of the amino acid in *i*-4 position. Remarkably, in the *C. velia* CLH a bulky Lys is located at *i*-4, which is expected to prevent the proposed mechanism (see Fig. 6). To illustrate the point in a more quantitative manner, a plot of the excitonic coupling between *a*403 and *a*406 for a range of LHC proteins is given in Fig. 7f. For this analysis, all pigments were parametrized as Chl *a*. Despite the data scattering, an overall trend towards a stronger coupling for the Asn ligand is present. Relating the excitonic coupling to the volume of the *i*-4 residue sidechain^58^ suggests a trend towards a decreasing coupling with an increasing volume (Fig. 7g), even for the His-coordinated systems. This can originate from the capacity of His to form the hydrogen bond with the carbonyl group in position 13^1^ of Chl *a*406 (see LHCII in Fig. 7b). Such an H-bond would be influenced as well by the size of the residue at position *i*-4. In order to verify this model, a mutation analysis of these positions, in analogy with previous studies^39,59^, would be highly informative.

FCP stands out from the trend outlined in Fig. 7g. However, this is the only LHC in which the chlorophyll at the *a*403 (*a*603) position is natively a Chl *c*, hence the simple dipole/site energy parametrisation to Chl *a* while retaining the Chl *c* geometry might not be fully adequate. However, if the estimated large coupling is correct, the Asn for His replacement in such a system assembled with only Chl *a* would lead to the formation of a red-shifted antenna complex. Remarkably, the diatom Lhcf15 also contains the Gly at the *i*-4, forming a GxxxN binding motif. This protein has been identified as the origin of the red-shifted antenna in *P. tricornutum*^37,38^. Lhcf15 thus fits the proposed mechanism, should *c*403 be occupied by Chl *a*. Pending further study, we note that this agrees with the (previously unexplained) elevated content of Chl *a* relative to Chl *c* in Lhcf15 antenna compared to FCP^60^.

The third representative of a far-red antenna is the redCLH of the alveolate *C. velia* formed by the proteins 1646_Red-CLH-1 and 532_Red-CLH-2 (Fig. 6). These proteins do not follow the present pattern since they possess a VCP/FCP-like binding motif [I/V]xxxH, but according to the phylogenetic analysis these proteins belong to a different LHC class than rVCP (Fig. S4), and more data are required to investigate the origin of the low-energy states in these LHCs.

### Concluding remarks

In the present work, we located the site of the red-shifted pigments of the eustigmatophyte antenna to the conserved core of the LHC protein. The primary factor for the red-shifted light adaptation appears to be a single amino acid residue exchange (Asn for His) of the Chl *a* ligand, supported by the presence of a small-sidechain residue at *i*-4 position. The proposed [A/G]xxxN motif in the helix B of LHC proteins thus emerges as a marker of a red-shifted antenna complex and can potentially serve as a genomic marker of the physiology adapted to the survival in shaded environments, in particular when present in an antenna complex not associated with photosystem I.

A yet unresolved general issue is the extent of the contribution of the tertiary and quaternary structure to direct pigment-pigment interactions. Resolving this in the red-shifted LHC complexes might bring important insights into other unsolved problems of the structure-function relationship in LHCs, such as the still elusive mechanisms of switching between the light-harvesting and the photoprotective conformations.

In perspective, understanding the factors governing the spectral tuning of LHCs could lead to the rational design of optimized light-harvesting systems for industrial cultivation of algae, without need of introducing the metabolic pathways necessary to produce red-shifted chlorophylls *d* or *f*.

## Experimental Methods

### Sample preparation

Cells of *Trachydiscus minutus* CCALA 838 were used as a source material for obtaining rVCP samples. *T. minutus* was batch-cultured in 5 L Erlenmeyer flasks in a freshwater WC medium^61^ at 20 °C. The cell cultures were stirred and bubbled with filtered air. Illumination was provided by a common halogen light bulb as a red-enhanced light source (intensity of 20 µmol photons m^−2^ s^−1^), following a rectangular wave cycle of 15 h light and 9 h dark^37^. rVCP was purified from solubilised thylakoid membranes^24^ by a combination of sucrose gradient centrifugation (0.1-1.1 M, 100,000 × g, 17 h) and size exclusion chromatography (Superdex 200 10/300 GL (GE Healthcare)) as previously described^34^.

### Transcriptome sequencing and assembly

*T. minutus* CCALA 838 was cultivated in liquid Bold’s basal medium (BBM)^62^. Total RNA was isolated using TRI Reagent® (TR 118) (Molecular Research Center, Inc., Cincinnati, USA), following standard procedures. Transcriptome sequencing was performed by the Institute of Applied Biotechnologies a.s. (Olomouc, Czech Republic) with the Illumina NovaSeq 6000 platform and the pair-end sequencing strategy. The obtained sequence data (36,062,624 reads) were quality trimmed and adapter clipped with Trimmomatic v0.39^63^. *De novo* transcriptome assembly was performed using Trinity v2.1.1^64^ and protein sequences were predicted with TransDecoder v5.5.0^65^ (https://github.com/TransDecoder/TransDecoder).

### Protein identification by MS/MS

Tandem mass spectroscopy (MS/MS) was used to identify the protein composition of purified rVCP^33^. In short, the samples were first analysed by denaturing polyacrylamide gel electrophoresis stained with Coomassie Blue, visible bands were excised from the gel, digested with trypsin, and analysed on a nano-scale UPLC coupled online to an ESI-Q TOF Premier Mass spectrometer (Waters, USA). Raw MS/MS data were processed and resulting peptides were subjected to a database search using PLGS2.3 software (Waters) against a custom database of *T. minutus* LHC protein sequences recovered from the *T. minutus* CCALA 838 transcriptome-derived protein sequence set using a blastp search (with *N. oceanica* VCP Lhcv1 sequence^33^ as a query).

### Sequence analysis

Comparison of LHCs amino acid sequences from *Pt* (*Pt*|Lhcf4, uniProt: B7FRW2; *Pt*|Lhcf15, uniProt: B7G8Q1)^44,45^ and *Tm* (DN2982, DN29098, and DN6201, inferred from the contigs TRINITY_DN2982_c0_g1_i1, TRINITY_DN29098_c0_g1_i1, and TRINITY_DN6201_c1_g1_i1, respectively, from the transcriptome assembly *de novo* generated for *T. minutus*, see above) was carried out by means of a multiple sequence alignment built using MAFFT (version 7)^66^. For phylogenetic analysis a set of LHC sequences was gathered by homology searches and literature survey. Specifically, candidate LHC sequences encoded by *T. minutus* were identified by searching the sequence set predicted from the *de novo* generated transcriptome assembly with hmmsearch (HMMER 3.0 package)^67^ as a query using a profile HMMM derived from the seed alignment of the Pfam protein family PF00504 (“Chlorophyll A-B binding protein”). Hits above the inclusion threshold were evaluated by blastp searches against the NCBI non-redundant protein sequence database and only evident members of the LHC family retained (excluding the divergent LHC-like LIL1 type). Some of the sequences proved truncated due to incomplete assembly of the respective transcripts. Virtually all of them could be completed by manually joining two or more separate contigs exhibiting perfect or near-perfect overlaps. The assembly was aided by an unpublished partial genome assembly and a separate alternative transcriptome assembly. The previously published sets of LHC sequences (excluding LIL1) encoded by the *Nannochloropsis oceanica* and *Microchloropsis* (=*Nannochloropsis*) *gaditana* genomes^33,68^ were retrieved from the respective databases and further refined by identification of additional, previously missed LHC family members and replacing some of the incorrect gene models with accurate protein sequences derived from transcriptome assemblies. The eustigmatophyte LHC sequences (with technical details listed in Supplementary Table S1) were combined with previously identified LHC sequences from the diatom *P. tricornutum*^69^ and a subset of LHC sequences from the alveolate *C. velia*, red algae (*Cyanidioschyzon merolae*, *Galdieria sulphuraria*, and *Porphyridium purpureum*), the green algal *Chlamydomonas reinhardtii*, and the plant *Arabidopsis thaliana*. The sequences were aligned with hmmalign (HMMER package) using the PF00504-derived profile HMM as the template and the --trimm option to remove the unaligned flanking (non-conserved) regions. The phylogenetic reconstruction was performed using the ETE3 3.1.2 pipeline^70^ as implemented on the GenomeNet (https://www.genome.jp/tools/ete/). Columns with more than twenty percent of gaps were removed from the alignment using trimAl v1.4.rev6^71^ and a maximum likelihood tree was inferred using IQ-TREE 1.5.5 ran with ModelFinder and tree reconstruction^72^, with PMB+F+R5 selected as the best-fit substitution model according to BIC. Branch support was tested by SH-like aLRT with 1000 replicates. For presentation purposes, the tree was visualised and adjusted using iTOL^73^.

### Optically Detected Magnetic Resonance

ODMR spectra were acquired in a home-built set-up described in detail previously^46,74^. Briefly, the light from a halogen lamp (250 W, Philips) is focused on the sample cell, which is immersed in a bath helium cryostat (all measurements were carried out at a temperature of 1.8 K), after being filtered through either a 5 cm CuSO_4_ solution (FDMR spectra) or a 10 cm water filter (triplet - minus-singlet, T-S, absorption-detected spectra). In FDMR experiments, the fluorescence is detected through bandpass filters (characterised by a full width at half maximum of about 10 nm) using a photodiode placed at 90° with respect to the excitation light direction, while in absorption-detected experiments, the light transmittance is detected with straight geometry through a monochromator (Jobin Yvon, mod. HR250). By sweeping the microwave frequency (MW source HP8559b, sweep oscillator equipped with a HP83522s plug-in and amplified by a TWT Sco-Nucletudes mod 10-46-30 amplifier) while detecting the fluorescence changes at specific wavelengths, the resonance transitions between spin sublevels of the triplet states can be determined. The microwaves are on/off amplitude modulated for selective amplification and the signal from the detector is demodulated and amplified using a Lock-In amplifier (EG&G, mod 5210). The microwave resonator, where the sample cell is inserted, consists of a slow pitch helix. FDMR spectra are presented as ΔF/F versus microwave frequency, where ΔF is the fluorescence change induced by the resonant microwave field and F is the steady-state fluorescence.

Microwave-induced T−S spectra can be collected by fixing the microwave frequency at a resonant value and sweeping the absorption detection wavelength. Compared to optical time-resolved absorbance spectroscopy on the triplet state, the ODMR technique allows selection (by the resonant microwave field) of specific triplet populations present in the sample, and in this way well resolved T-S spectra associated with specific chromophores can be obtained.

### Pulse and time-resolved EPR

The rVCP samples were concentrated to a concentration of about 350 μg/ml of Chl *a*. Glycerol, previously degassed by several cycles of freezing and pumping, was added (60% v/v) just before freezing to obtain a transparent matrix. The sample of Chl *a* in Triton X-100 micelles was obtained by adding a few microliters of a concentrated solution of the pigment (SIGMA) dissolved in methanol to 1 ml of 1 mM Triton X-100, as previously reported^49^.

Pulse EPR experiments were performed on a Bruker ELEXSYS E580 spectrometer, equipped with a dielectric cavity (Bruker ER 4117-DI5, TE_011_ mode), an Oxford CF935 liquid helium flow cryostat, and an Oxford ITC4 temperature controller. The microwave frequency was measured using a frequency counter (HP5342A). The temperature was controlled in a helium-flow and all experiments were conducted at 50 K, disabling magnetic field modulation and using pulsed sample photo-excitation from a Nd:YAG pulsed laser (Quantel Brilliant) equipped with second and third harmonic modules and an optical parametric oscillator (OPOTECH) (λ=440 nm, pulse length = 5 ns, E/pulse ≅ 1.5 mJ, 10 Hz repetition time). The pulse EPR experiments were recorded using a two-pulse Electron Spin Echo (ESE) sequence (flash-delay after flash-π/2-τ-π-τ-echo), where the echo intensity was registered as a function of the magnetic field. The microwave π/2-pulse was of 16 ns and the delay τ was set at 300 ns.

The TR-EPR spectra were performed on the same instrument, where the EPR direct-detected signal was recorded with a LeCroy 9300 digital oscilloscope, triggered by the laser pulse. The temperature was controlled in a nitrogen-flow and all experiments were conducted at 50 K and 120 K. For every field position, 300 transient signals were averaged. To eliminate the laser background signal, transients accumulated at off-resonance field positions were subtracted from those on resonance.

Simulations of the powder spin-polarised triplet spectra were performed using a program written in Matlab®, with the aid of the EasySpin routine (ver. 5.2.25)^75^, based on the full diagonalisation of the triplet state spin Hamiltonian, including the Zeeman and electron-electron magnetic dipole interactions, considering a powder distribution of molecular orientations with respect to the magnetic field direction. Input parameters are the triplet state sublevel populations, the zfs parameters, the linewidth, and the isotropic g value.

Calculations of the sublevel triplet state populations of the acceptor (Car), starting from those of the donor (Chl), were performed using a home-written program in Matlab® software previously described^6,42,52^, utilising the X-ray coordinates of *Pt* FCP^10^.

## Data availability

Raw sequencing reads and the transcriptome assembly from *Trachydiscus minutus* CAUP 838 are available as NCBI BioProject PRJNA1078143. Protein sequences inferred from the *T. minutus* transcriptome assembly are available from Figshare (DOI: 10.6084/m9.figshare.25706553). The authors declare that the data supporting the findings of this study are available from the corresponding authors upon request.

## Acknowledgments

The skilled technical assistance of Ivana Hunalová, František Matoušek, and Sabrina Mattiolo is gratefully acknowledged. The Czech Science Foundation (20-01159S to D. Bína and 21-19664S to M.E.), the MEMOVA project (EU Operational Programme Research, Development and Education No. CZ.02.2.69/0.0/0.0/18_053/0016982), the University of Padova (P-DiSC#02BIRD2020-UNIPD to M.B.), and the MUR (support to D.C and A.A. under the National Recovery and Resilience Plan (NRRP/PNRR), Mission 4, Component C2, Investment 1.1, Call for tender No. 104 of the 02/02/2022 by the Italian Ministry of University and Research (MUR), funded by the European Union – NextGenerationEU– Project Title “Extending the red limit of oxygenic photosynthesis: basic principles and implications for future applications (20224HJWMH)” – CUP C53D23004620006 Grant Assignment Decree No. 1017 adopted on 07/07/2023 by MUR) financially supported this work. Institutional support RVO: 60077344 is also acknowledged.

## Author Contributions

A.A., D. Bína, and R.L. conceived and designed the research and coordinated the project. D. Bína extracted and purified rVCP and analysed its composition by tandem mass spectrometry. A.A., M.B., and D.C. performed the EPR and ODMR experiments and analysed the data. D. Barcytė and M.E. obtained the *T. minutus* transcriptome. A.A., D. Bína, D. Barcytė, and M.E. analysed the LHCs polypeptide sequences. A.A. wrote the original draft of the paper. All authors reviewed and edited the manuscript.

