## Supplementary material for "Origin of the far-red absorbance in eustigmatophyte algae red-shifted Violaxanthin-Chlorophyll *a* Protein": Figures S1-S4 and Tables S1 and S2


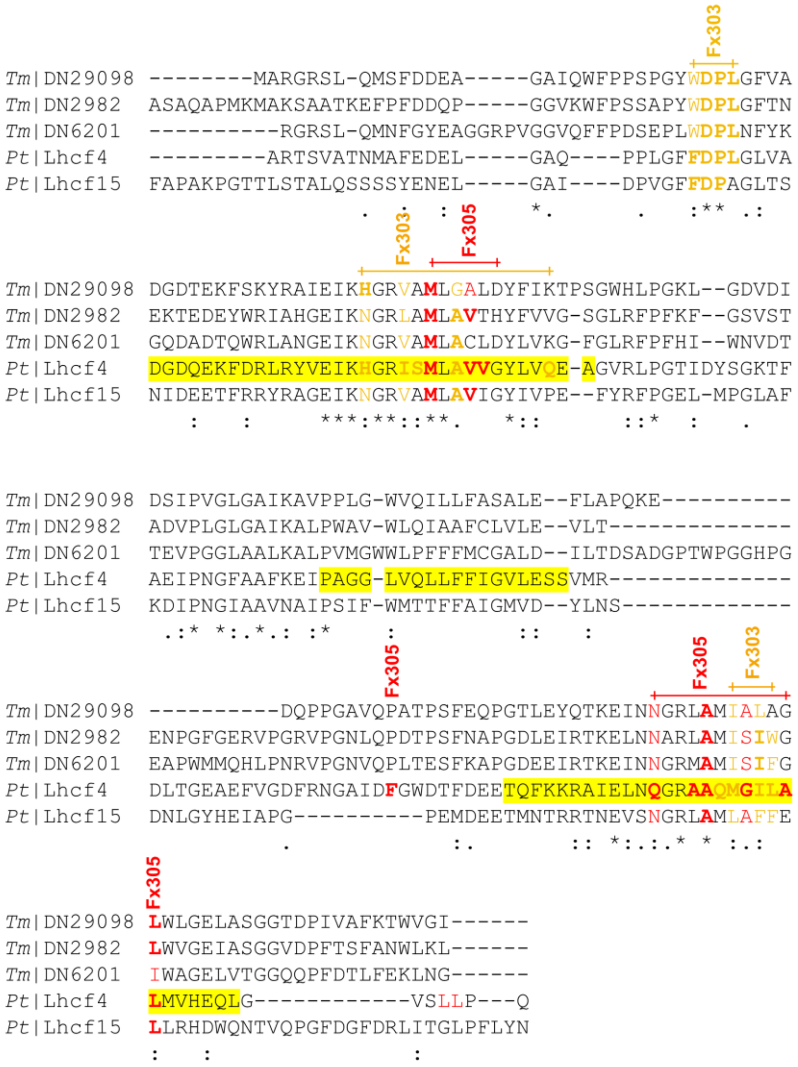


**Fig. S1| Alignment of polypeptide sequences of LHCs.** Polypeptide sequence alignment of the LHCs from *T. minutus* (*Tm*) and *P. tricornutum* (*Pt*)[44,45] were used for the alignment. Yellow background, transmembrane helix (derived from the crystallographic structures 6A2W[10]). Residues in a 4 Å radius from the carotenoids are color-coded: orange, Fx303; red, Fx305.

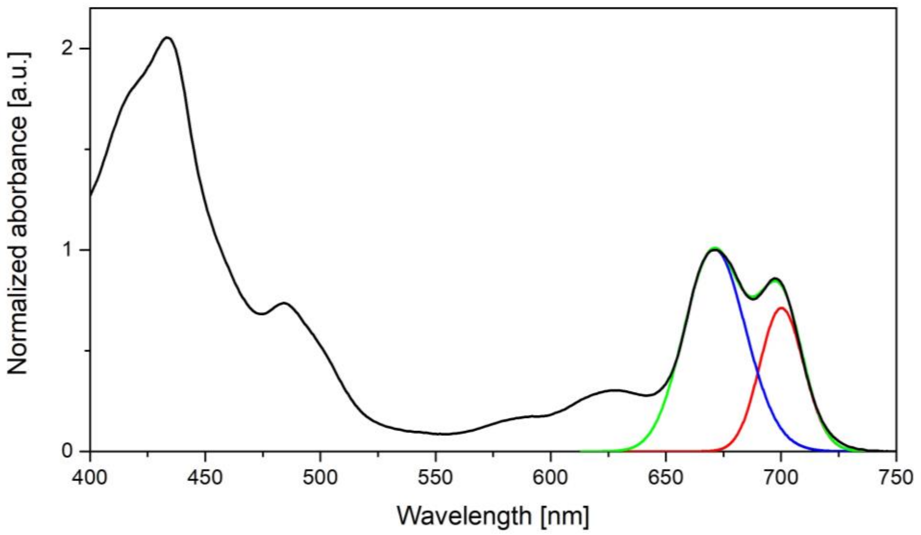


**Fig. S2| Room temperature absorption spectrum of *T. minutus* rVCP (black), with a spectral deconvolution of the $Q_y$ region as a sum (green) of two Gaussian contributions (blue and red)**. The parameters of the two Gaussians components are: peak position (671.0 and 700.2 nm), FWHM (696.4 and 456.7 $cm^{-1}$), and their area expressed as a percentage (68.3 and 31.7 %).

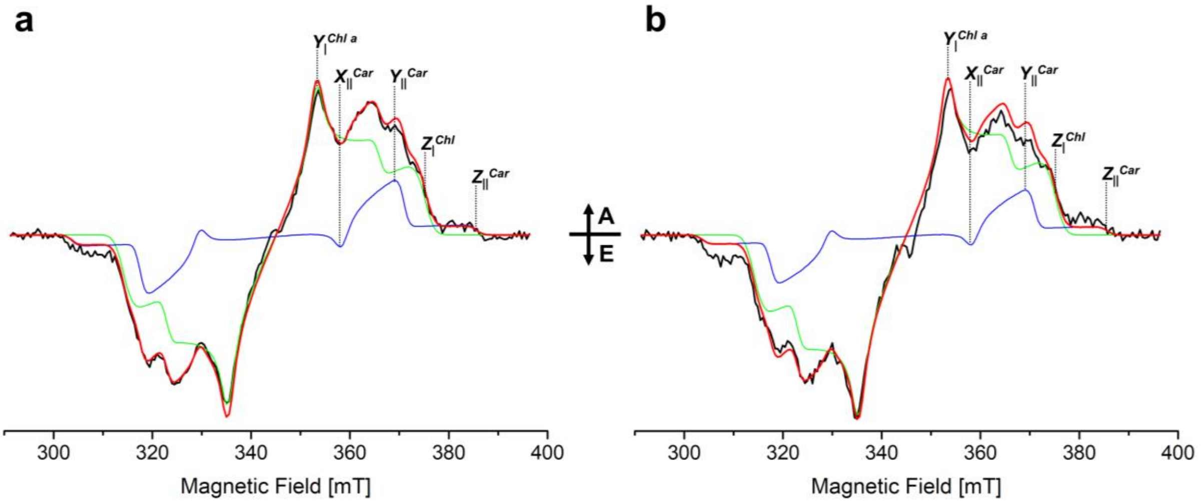


**Fig. S3| TR-EPR spectra at 700 ns after the laser pulse of rVCP (black lines) detected at (a) 50 K and (b) 120 K.** Reconstitutions of the FCPs TR-EPR spectra (red lines) have been obtained from the sum of $^3$Car (blue lines) and $^3$Chl *a* (green lines) contributions, weighted as reported in the figure. The polarizations of the simulated $^3$Car components was determined on the basis of atomic coordinates for the acceptor-donor pairs derived from the crystallographic structure[10] and an initial donor $^3$Chl polarisation ($P_x$:$P_y$:$P_z$ = 0.375:0.425:0.20)[5], resulting in a $^3$Car for both Fx303 and Fx305 (see Fig. 6). The simulated $^3$Car spectra were calculated using the following parameters: D = -41.0 mT; E = -4.1 mT; triplet polarization ($P_x$:$P_y$:$P_z$ = 0.41:0.20:0.39); linewidths ($lw_x$, $lw_y$, $lw_z$) = (2.0, 2.0, 2.5) mT. The simulated $^3$Chl *a* spectra were calculated using the following parameters: D = 30.5 mT; E = -4.2 mT; triplet polarization ($P_x$:$P_y$:$P_z$ = 0.375:0.425:0.20); linewidths ($lw_x$, $lw_y$, $lw_z$) = (2.0, 2.0, 3.5) mT. Canonical transitions have been highlighted on the high-field half of the spectra. A = absorption, E = emission.

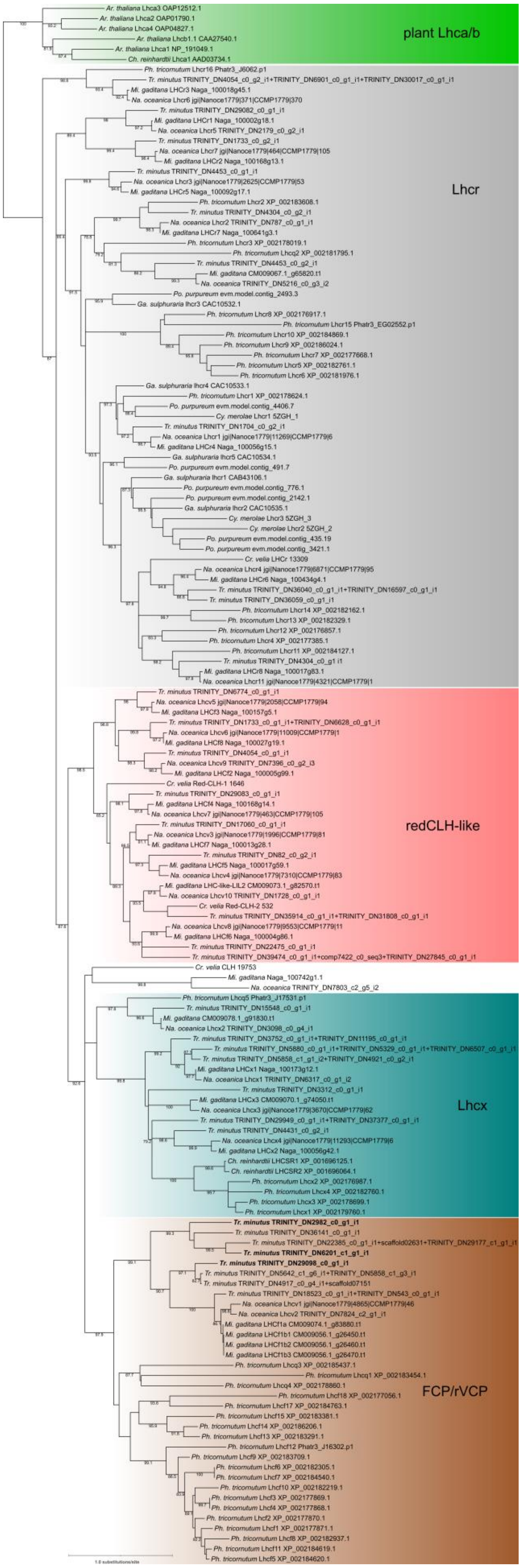


**Fig. S4| Maximum likelihood phylogenetic tree of LHC sequences for the purpose of classification of the *Trachydiscus minutus* (*Tm*) antenna sequences**. The tree was inferred using IQ-TREE from a multiple alignment of protein sequences (172 amino acid positions) using the substitution model PMB+F+R5. The numbers at branches correspond to support values calculated with SH-like aLRT (1000 replicates); displayed are values ≥75. The tree is rooted arbitrarily, the major subgroups of LHC proteins are delimited with a different colour background. Included were LHC sequences identified in genome or transcriptome assemblies of the eustigmatophytes *T. minutus* CCALA 838, *Nannochloropsis oceanica*, and *Microchloropsis* (=*Nannochloropsis*) *gaditana* and the diatom *Phaeodactylum tricornutum*, plus selected sequences from *Chromera velia* (Alveolata), *Cyanidioschyzon merolae*, *Galdieria sulphuraria*, and *Porphyridium purpureum* (Rhodophyta), *Chlamydomonas reinhardtii* (Chlorophyta), and *Arabidopsis thaliana* (Embryophyta). Gene names are provided following the previous literature (we note that the nomenclature is highly inconsistent). The labels given in bold (in the FCP/rVCP group) correspond to proteins that were found to constitute the *T. minutus* rVCP complex using MS/MS.

**Table S1**| MS/MS identification peptides from proteins forming the rVCP complex of *T. minutus*

| protein | mass [kDa] | peptides | start | end | score | coverage [%] |
|---|---|---|---|---|---|---|
| **DN2982** | 22.2 | (K)EFPFDDQPGGVK | 26 | 37 | 90.142 | 46 |
| | | (K)FGSVSTADVPLGLGAIK | 94 | 110 | 240.52 | |
| | | (R)IAHGEIK | 63 | 69 | 92.319 | |
| | | (R)LAMLAVTHYFVVGSGLR | 73 | 89 | 185.9 | |
| | | (K)TEDEYWR | 56 | 62 | 96.51 | |
| | | (R)VPGNLQPDTPSFNAPGDLEIR | 144 | 164 | 173.15 | |
| | | (K)WFPSSAPYWDPLGFTNEK | 38 | 55 | 184.65 | |
| | | (K)WFPSSAPYWDPLGFTNEKTEDEYWR | 38 | 62 | 117.9 | |
| **DN29098** | 24.6 | (K)AVPPLGWVQILLFASALEFLAPQK | 143 | 166 | 37.769 | 45 |
| | | (K)EDQPPGAVQPATPSFEQPGTLEYQTK | 167 | 192 | 302.02 | |
| | | (R)LAMIALAGLWLGELASGGTDPIVAFK | 199 | 224 | 218.79 | |
| | | (K)LGDVDIDSIPVGLGAIK | 126 | 142 | 244.71 | |
| | | (K)TPSGWHLPGK | 116 | 125 | 103.8 | |
| | | (R)VAMLGALDYFIK | 104 | 115 | 121.8 | |
| **DN6201** | 24.9 | (K)APGDEEIR | 182 | 189 | 69.905 | 43 |
| | | (R)FPFHIWNVDTTEVPGGLAALK | 100 | 120 | 171.51 | |
| | | (K)GQDADTQWR | 64 | 72 | 146.03 | |
| | | (R)MAMISIFGIWAGELVTGGQQPFDTLFEK | 198 | 225 | 178.05 | |
| | | (R)PVGGVQFFPDSEPLWDPLNFYK | 42 | 63 | 155.61 | |
| | | (R)VPGNVQPLTESFK | 169 | 181 | 200.76 | |

**Table S2**| Global fitting parameters used for the reconstruction of the $^3$Chl FDMR spectra in Fig. 3b.

| | **D-E** [MHz] | **D+E** [MHz] | **FWHM** [MHz] | **D** [$cm^{-1}$] | **E** [$cm^{-1}$] |
|---|---|---|---|---|---|
| **$^3Chl_1$** | 726 ± 1 | 951 ± 1 | 31 ± 1 | 0.02792 ± 0.00001 | 0.00375 ± 0.00001 |
| **$^3Chl_2$** | 734 ± 1 | 987 ± 1 | 31 ± 1 | 0.02865 ± 0.00001 | 0.00421 ± 0.00001 |
