## Supplementary figures and images for "Origin of the far-red absorbance in eustigmatophyte algae red-shifted Violaxanthin-Chlorophyll *a* Protein"

### Figure S4

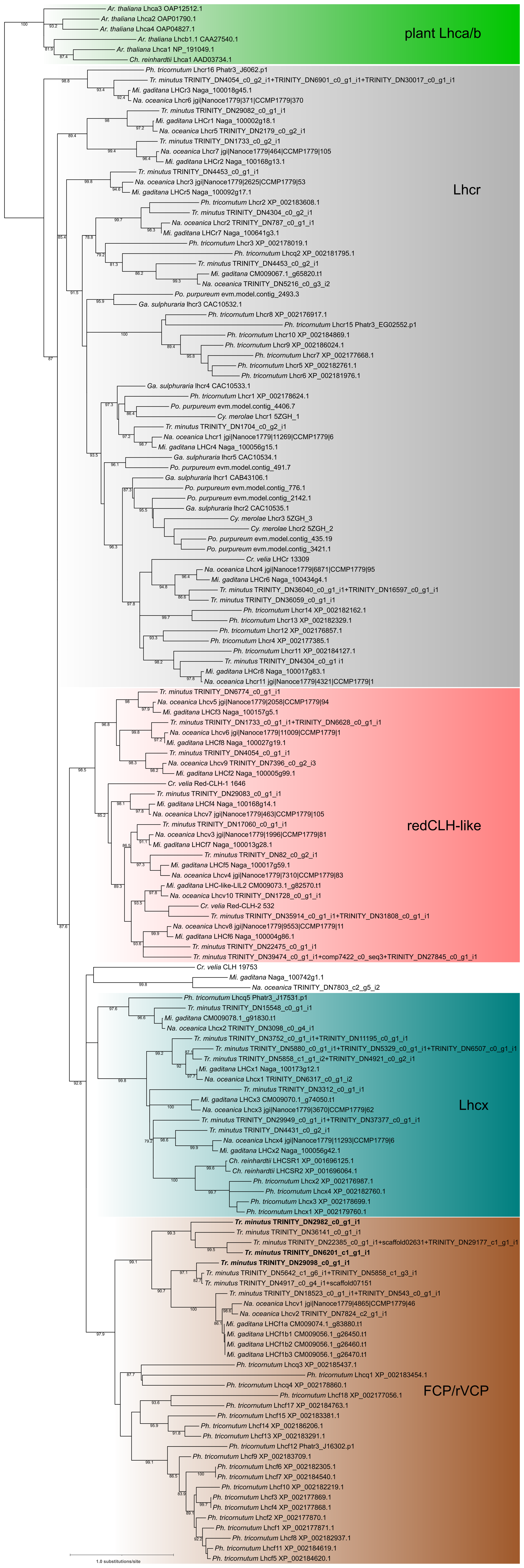
